# RNA targeting with CRISPR-Cas13a facilitates bacteriophage genome engineering

**DOI:** 10.1101/2022.02.14.480438

**Authors:** Jingwen Guan, Agnès Oromí-Bosch, Senén D. Mendoza, Shweta Karambelkar, Joel Berry, Joseph Bondy-Denomy

**Author notes:** Department of Biology, Massachusetts Institute of Technology, Cambridge, MA, USA.

## Abstract

The viruses that infect bacteria, bacteriophages (or phages), possess numerous genes of unknown function. Genetic tools are required to understand their biology and enhance their efficacy as antimicrobials. *Pseudomonas aeruginosa* jumbo phage ΦKZ and its relatives are a broad host range phage family that assemble a proteinaceous “phage nucleus” structure during infection. Due to the phage nucleus, DNA-targeting CRISPR-Cas is ineffective against this phage and thus there are currently no reverse genetic tools for this family. Here, we develop a DNA phage genome editing technology using the RNA-targeting CRISPR-Cas13a enzyme as a selection tool, an anti-CRISPR gene (*acrVIA1*) as a selectable marker, and homologous recombination. Precise insertion of foreign genes, gene deletions, and the addition of chromosomal fluorescent tags into the ΦKZ genome were achieved. Deletion of *phuZ*, which encodes a tubulin-like protein that centers the phage nucleus during infection, led to the mispositioning of the phage nucleus but surprisingly had no impact on phage replication, despite a proposed role in capsid trafficking. A chromosomal fluorescent tag placed on gp93, a proposed “inner body” protein in the phage head revealed a protein that is injected with the phage genome, localizes with the maturing phage nucleus, and is massively synthesized around the phage nucleus late in infection. Successful editing of two other phages that resist DNA-targeting CRISPR-Cas systems [OMKO1 (ΦKZ-like) and PaMx41] demonstrates the flexibility of this method. RNA-targeting Cas13a system holds great promise for becoming a universal genetic editing tool for intractable phages. This phage genetic engineering platform enables the systematic study of phage genes of unknown function and the precise modification of phages for use in a variety of applications.

**Graphical abstract:** 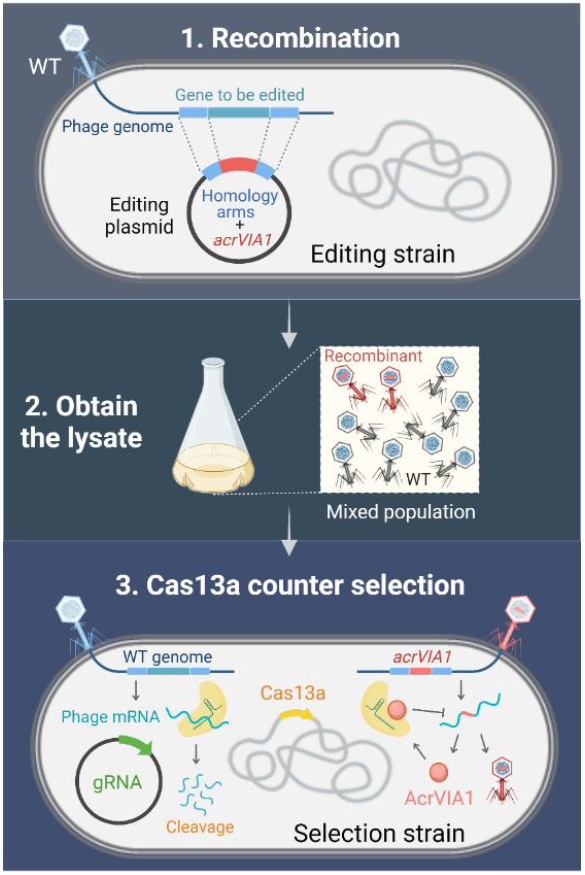

## Introduction

Bacteriophages are viruses that infect bacteria and can cause their lysis after replication. In recent decades, the rapid emergence of multi-antibiotic resistant bacterial pathogens and simultaneous decline in the discovery of new antibiotics has rekindled interest in the use of phages as alternative antimicrobial therapeutics (phage therapy) [1, 2]. Phages offer many advantages over antibiotics, including high specificity and efficient self-propagation in the presence of their bacterial host [3–5]. However, host range limitations and the rapid emergence of phage resistance in clinical strains present barriers for furthering phage therapy [1, 3, 6]. Phage genome engineering may help overcome these hurdles [7, 8]. Robust phage engineering tools can aid fundamental discoveries, broaden host range, enhance evasion of host antiviral defense systems, and reduce phage toxicity and immunogenicity [9–12]. Phage engineering techniques often utilize homologous recombination (HR) with a template plasmid [13, 14], coupled with a selective pressure such as CRISPR-Cas targeting. CRISPR-Cas systems (clustered regularly interspaced short palindromic repeats and CRISPR-associated proteins) are adaptive anti-phage immune systems in prokaryotes [15, 16]. CRISPR-Cas programmable targeting enables effective enrichment for phage recombinants by removing wild-type phages from the population and has been coupled with the integration of an anti-CRISPR gene as a selectable marker [17].

To date, all CRISPR-based screening tools used in phage engineering recognize and target phage genomic DNA. However, due to the everlasting evolutionary battle between bacteria and phages in nature, phages have amassed various strategies to circumvent DNA-targeting immunity, including anti-CRISPR proteins, DNA base modifications, and genome segregation [18, 19]. *P. aeruginosa* jumbo phage ΦKZ is resistant to a broad spectrum of DNA-targeting immune systems via assembly of a proteinaceous “phage nucleus” structure that shields phage DNA during replication [20, 21]. This phage family is thus a great candidate for use as a phage therapeutic and likely possesses other fascinating fundamental biology. However, no genetic tools are available for this phage family and most studies have relied on plasmid-based over-expression [22].

Although the phage nucleus prevents DNA-targeting, the mRNA-targeting CRISPR-Cas13a system (type VI-A) [23] effectively inhibits ΦKZ replication by degrading phage mRNA that is exported from the phage nucleus to the cytoplasm [20]. Here, we develop the CRISPR-Cas13a system as a novel genetic engineering approach for ΦKZ. Using Cas13a to target an essential transcript, we select for phage DNA that has undergone homologous recombination resulting in a desired genetic change along with the acquisition of an anti-Cas13a *trans* gene, *acrVIA1* (derived from Listeriophage ΦLS46 [24]), as a selectable marker. This approach allows us to precisely insert foreign gene fragments into the ΦKZ genome, knock out non-essential genes, and fuse fluorescent tags to individual genes. Importantly, the same guide can be used for any genomic manipulation of a single phage as engineered phages are identified based on the acquisition of the Cas13a inhibitor, not a change in the target sequence. Our work establishes a Cas13a-based phage engineering strategy that could be a universally powerful tool for engineering phages.

## Results

### Optimization of CRISPR-Cas13a for efficient phage targeting

Cas13a is an RNA-guided RNA nuclease that can block ΦKZ replication in *P. aeruginosa* PAO1. We previously targeted ΦKZ by expressing crRNA guides from a plasmid (Figure 1A, Version 1) [20]. For the effective elimination of WT phages in the population, we first sought to enhance activity of crRNA guides. We designed Version 2 (V2) with the repeat-spacer-repeat unit moved to the +1 transcription start site and the second direct repeat (DR) mutated to remove repeat homology (Figure 1A). To further stabilize the crRNA cassette, we omitted the second DR and generated V3 (Figure 1A), which could prevent recombination between the repeats. Using the same spacer, both V2 and V3 provided more robust defense against phage JBD30 and ΦKZ compared with V1 (Figure 1A). For simplicity, we selected the V3 cassette that has a single repeat to express crRNAs against ΦKZ. We designed multiple spacer sequences to target various ΦKZ gene transcripts. Strong targeting was observed for some crRNAs, to the point that escaper phages could be isolated, such as the two spacers matching *orf120* and *orf146* transcripts, but not all crRNAs were efficacious (Figure 1B and S1). Given the variability of targeting efficiency, for the remainder of this report, we use *orf120*-targeting gRNA#2 as our primary guide to screen for engineered phages. We refer the PAO1 strain simultaneously expressing Cas13a and *orf120-* gRNA#2 to as Cas13a counter-selection strain. We describe below how the same guide can be used to facilitate the engineering of distinct genomic loci.

**Figure 1.**
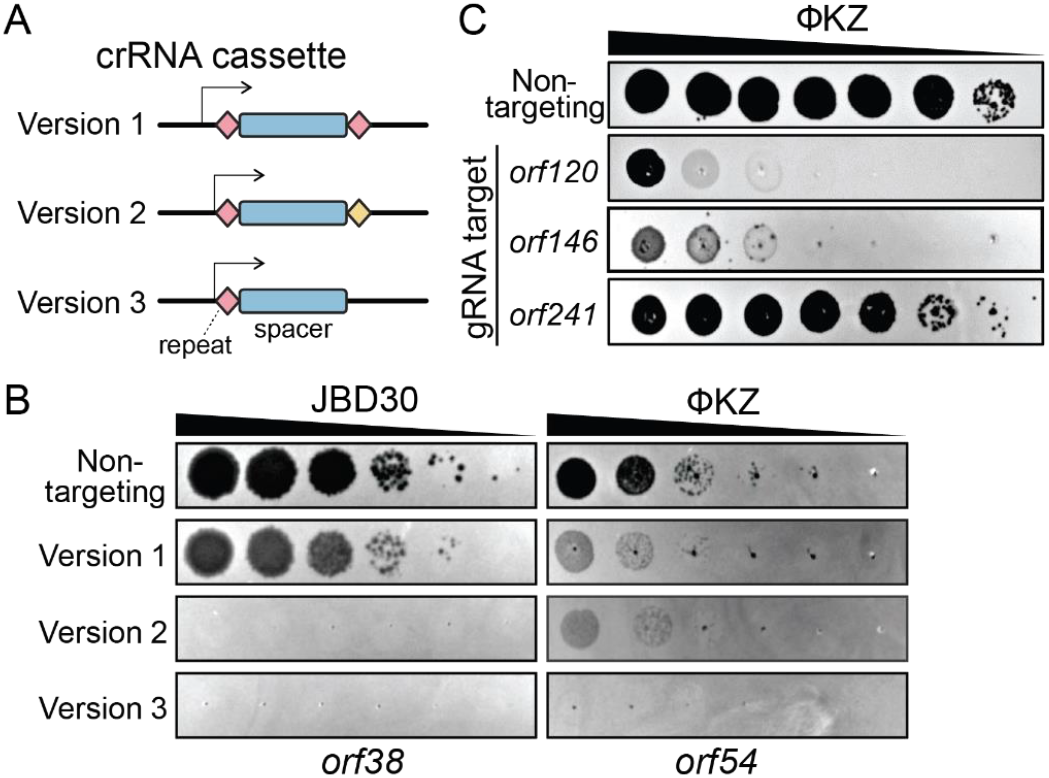
Optimization of CRISPR-Cas13a crRNA expression vector for efficient phage interference. (A) Schematic of three versions of CRISPR-Cas13a crRNA cassette (B) Efficiency of plaquing of three versions of crRNA cassette targeting two unrelated phage strains: JBD30 and ΦKZ. Wild-type phages were spotted in ten-fold serial dilutions (left to right) on a lawn of *P. aeruginosa* PAO1 expressing *Lse*Cas13a from the chromosome and harboring indicated crRNA expression vectors. The cassettes carried the same spacer sequences targeting transcripts of *orf38* of JBD30 and *orf54* of ΦKZ, respectively. (C) Cas13a interference efficiency of distinct crRNAs against ΦKZ using Version 3 crRNA expression vector. The most efficient crRNA that targets the *orf120* transcript was selected for genetic engineering of ΦKZ. All plaque assays were replicated three times yielding similar results.

### Isolation of ΦKZ recombinants by CRISPR-Cas13a counter selection

To avoid disrupting any essential genes that are required for phage replication, we first attempted to insert *acrVIA1* immediately downstream of ΦKZ major capsid gene (*orf120*). A template DNA substrate for homologous recombination, composed of ~600-bp homology arms flanking *acrVIA1* was cloned into a plasmid, referred to as an editing plasmid (Figure 2A). After infecting a PAO1 strain with the editing plasmid to allow recombination, the phage lysate was then titrated on a lawn of the Cas13a counter-selection strain to eliminate WT phages. To screen for recombinants, individual plaques were examined for *acrVIA1* integration via PCR, showing that 8 out of the 16 tested phage plaques generated the expected 1.6 kbp band, which was not detectable in a WT plaque (Figure 2B). Amplification of the entire region also revealed the expected size increase, from ~1.7 kbp of WT to ~2.5 kbp of recombinants, and Sanger sequencing confirmed the correct integration junction (Figure 2C). Recombinant phages propagated well on hosts expressing crRNAs targeting other genomic loci due to the expression of *acrVIA1*, which abolishes Cas13a immunity regardless of the crRNA sequence (Figure 2D). Three randomly selected phages that escaped Cas13a targeting but screened negative for the *acrVIA1* integration contained genomic deletions ranging from 27 bp to 69 bp starting immediately downstream of the *orf120* stop codon, disrupting the protospacer (Figure S2). While the crRNA used is therefore not inescapable, the recombination efficiency to insert the selectable marker is clearly efficient enough to enable facile identification of the desired mutants.

**Figure 2.**
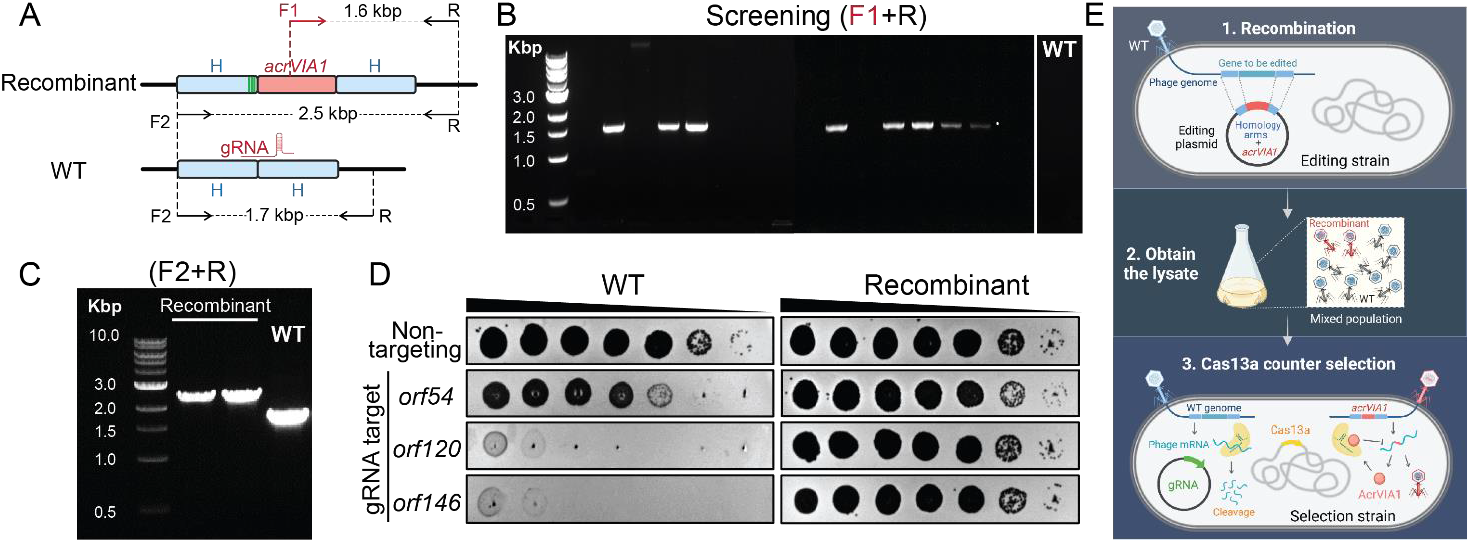
Screening for ΦKZ recombinants by CRISPR-Cas13a counter selection. (A) Schematic of WT ΦKZ and recombinant genomes at the editing site. The *acrVIA1* gene, shown as a red rectangle, was inserted downstream of *orf120*, with up- and downstream of homology arms (H) indicated by blue rectangles. Green stripes represent synonymous mutations that were introduced to the homology region to prevent crRNA targeting for recombinant phages. F and R indicate forward and reverse primers, respectively, being used to confirm the insertion of *acrVIA1*. (B) Recombinant phages were screened by PCR using primer F1 and R. (C) Recombinant phage plaques underwent 3 rounds of purification and were further confirmed by PCR using primer F2 and R. (D) Plaque assays showing robust anti-CRISPR activities acquired by recombinant ΦKZ against distinct crRNAs, owing to the successful expression and execution of the incorporated AcrVIA1. (E) Workflow of phage genome engineering using CRISPR-Cas13a. The strategy can be divided into three steps. In the first step, an editing plasmid is constructed to introduce desired genetic modifications, such as insertion, deletion, and tag-fusion, and *acrVIA1* flanked by up- and downstream homology arms matching the phage genome. This plasmid is transformed into a bacterial host strain, referred to as the editing strain, followed by infection by wild-type phages. In the second step, the infection culture is harvested, yielding a mixed phage lysate, containing WT phages, recombinants, and escaper mutants. In the last step, the lysate is plated on the selection strain harboring Cas13a on the bacterial chromosome and the most effective gRNA targeting WT phages. Recombinants and escaper mutants will be recovered to form visible plaques on the cell lawn, while WT phages are eliminated by CRISPR-Cas13a via cleavage of targeted phage mRNA. Recombinants are screened by PCR using appropriate primers and further confirmed by sequencing. Remarkably, AcrVIA1 produced by recombinant phages enables ineffectiveness of Cas13a regardless of any gRNA. Accordingly, the same guide can be used for any genomic manipulation of the same phage strain, greatly reducing the effort on seeking for efficient gRNAs to edit distinct genomic loci.

To test the flexibility of this nascent genetic technology (Figure 2E) and generate new biological insights of phage ΦKZ, we next knocked out (or attempted to knock out) multiple genes; *phuZ (orf39), orf54, orf89-orf93, orf93, orf146, orf241*, and *orf241-orf242* (summarized in Table 1), in addition to attempting to add chromosomal fluorescent tags onto *orf54* and *orf93*. The successes, failures, and new insights gained are discussed below. Whole genome sequencing of two deletion mutants, Δ*phuZ* and Δ*orf93*, revealed no other mutations. The accuracy of this system highlights an important advantage of adapting an RNA-targeting system to select for edits in DNA phage genomes, compared to direct DNA cleavage.

**Table 1.** Summary of phage mutants engineered by CRISPR-Cas13a.

| Phage | Gene No. /Genomic site | Identified protein | Modification | Plaque#<sup>1</sup> | screening% | Isolated or not? |
| --- | --- | --- | --- | --- | --- | --- |
| **ΦKZ** | Upstream of *orf54* | - | Insertion | 17 | 41.2% | Yes |
|  | Downstream of *orf120* | - | Insertion | 16 | 50.0% |  |
|  | *orf39* | PhuZ | Deletion | 17 | 52.9% |  |
|  | *orf93* | Inner body protein | Deletion | 20 | 20.0% |  |
|  |  |  | gp93-mNeonGreen fusion | 22 | 18.2% |  |
|  | *orf241* | Hypothetical | Deletion | 23 | 26.1% |  |
|  | *orf241, orf242* | Hypothetical | Double deletions | 12 | 41.7% |  |
|  | *orf54* | Shell | Deletion | 15 | 20.0% | No |
|  |  |  | mCherry-gp54 fusion | 24 | 8.3% |  |
|  |  |  | gp54-mCherry fusion | 11 | 9.1% |  |
|  |  |  | gfp11-gp54 fusion | 57 | 7.0% |  |
|  | *orf89, orf90, orf91, orf92, orf93* | Head structural proteins | Multiple deletions | 17 | 11.8% |  |
|  | *orf146* | Structural protein | Deletion | 86 | 0.0% |  |
| **OMKO1** | Downstream of the capsid gene | - | Insertion with a barcode | 24 | 70.8% | Yes |
|  | Upstream of the shell gene | - | Insertion | 12 | 50.0% |  |
| **PaMx41** | *orf24* | Hypothetical | Deletion | 8 | 100.0% | Yes |

<sup>1</sup>Number of plaques analyzed by PCR to screen for recombinants.

### Characterization of PhuZ and gp93 using engineered ΦKZ mutants

PhuZ (gp39) is a tubulin homolog conserved across “group 1” jumbo phages and some megaphages [25, 26]. It assembles a bipolar spindle to center the phage nucleus during phage intracellular development [27, 28], and “treadmill” newly synthesized phage capsids from the cell inner membrane to the phage nucleus for DNA packaging [29]. These functions made us speculate that PhuZ might be essential for phage growth, however, this is not the case. Δ*phuZ* mutants exhibited a similar burst size (24 phage particles per infected bacterial cell *vs*. 19 of WT phage) under our experimental conditions. While cells infected with WT phages or Δ*phuZ* mutants complemented *in trans* had phage nuclei in the center of the cell ~80% of the time, the localization of the phage nucleus showed a wide distribution in cells infected by Δ*phuZ* mutants (Figure 3A-3B, Movie S1). This is consistent with the previous findings that *trans* over-expression of catalytic mutant PhuZ resulted in mispositioning of the phage nucleus [28, 30]. ~25% of mutant-infected cells still positioned the phage nucleus at the cell center (Figure 3B), a phenotype most commonly seen in shorter cells (*Pearson* correlation coefficient = 0.486, *p* < 0.001), in contrast with WT infection where no correlation was observed (*Pearson* correlation coefficient = 0.029, *p* = 0.505) (Figure 3C). Considering that PhuZ is proposed to traffic phage capsids from the cell inner membrane to the phage nucleus [29, 31], we speculate that this is only required under specific conditions or not at all. Taken together, these data suggested that PhuZ positions the phage nucleus at the cell center during infection, but the removal of PhuZ does not have a significant impact on ΦKZ growth in laboratory conditions.

**Figure 3.**
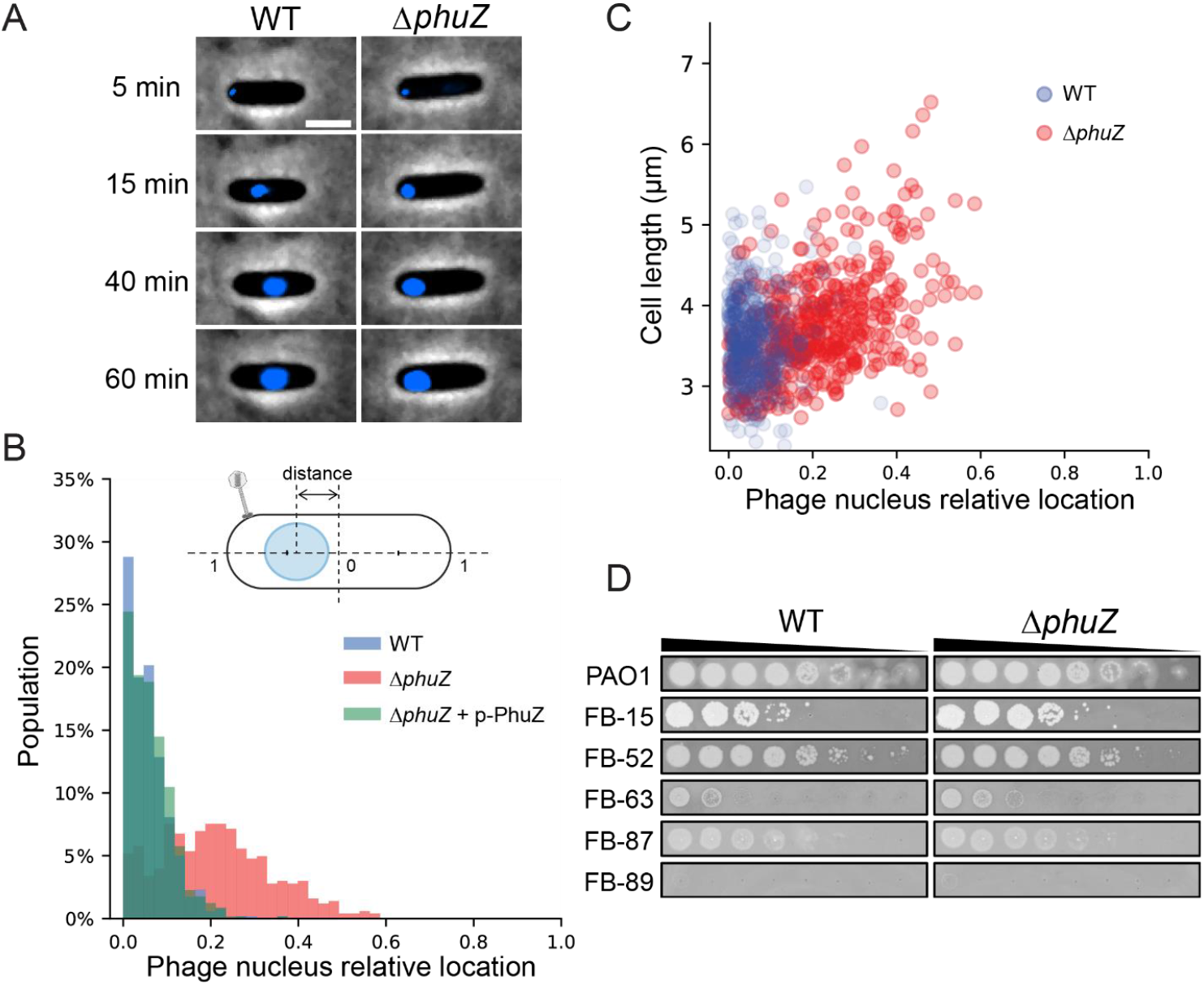
Absence of PhuZ causes mispositioning of the phage nucleus. (A) Left: a representative cell infected by WT ΦKZ. Right: a representative cell infected by Δ*phuZ* mutant. Phage DNA is stained by DAPI and shown as blue signals. The phage nucleus is mispositioned near the cell polar region upon infection by Δ*phuZ*, in contrast to the WT infection where the phage nucleus is centered. Scale bar denotes 2 μm. (B) Distribution of subcellular location of the phage nucleus in PAO1 cells infected by WT (blue, N = 521) and Δ*phuZ* (red, N = 503), and a PAO1 strain expressing wild-type PhuZ *in trans* (p-PhuZ) and infected by Δ*phuZ* (light green, N = 573). The diagram of an infected cell is shown on the top. The phage nucleus location is defined as the relative distance between the cell center and the nucleus center. (C) The phage nucleus location is plotted against the cell length for WT and Δ*phuZ*. The phage nucleus position in Δ*phuZ*-infected cells positively correlates with the cell length with a *Pearson* correlation coefficient of 0.486, *p* < 0.001, whereas there is no such correlation for WT infection with a *Pearson* correlation coefficient of 0.029, *p* = 0.505. (D) Efficiency of plaquing of WT and Δ*phuZ* on representative *P. aeruginosa* clinical strains. (The full panel of plaque assays is presented in Figure S3)

*orf93* encodes gp93, a high copy number “inner body (IB)” protein that is packaged in the phage head [32, 33]. Deletion of *orf93* also yielded viable phage with no obvious growth defect. We next analyzed an inserted fluorescent chromosomal label at the C-terminus of the protein, which is notable as the first chromosomal protein tag in this phage family. Labeled gp93 was observed in the mature virion (Figure 4A), as predicted by previous mass spectrometry studies [33]. Excitingly, time-lapse movies revealed the fluorescently labeled protein being injected with the phage DNA at the cell pole (Figure 4B, Movie S2) and subsequently translocating to the cell center where it remained bound to the phage nucleus. As new gp93-mNeonGreen was expressed from the phage genome, more and more green signals concentrated on the surface of the phage nucleus, while some foci appeared on the cell inner membrane. Finally, cells lysed and released fluorescent phage progeny. To confirm that the protein that appears to be injected was not rapidly synthesized *de novo*, we monitored the infection behaviors of WT phages loaded with gp93-mNeonGreen expressed from a plasmid during phage production, but where no new fluorescent protein could be made during infection (Figure 4C, Movie S3). Similar to the engineered phage, the phage particles were fluorescent, injected the labeled protein, and the green focus migrated from the cell pole to the cell center along with phage DNA on the surface of the phage nucleus until cell lysis. Therefore, the IB protein gp93 is not only packaged in the phage head, but may also play a role in phage development and maturation. The ability to chromosomally label phage proteins, as demonstrated here, will be beneficial for characterizing ΦKZ virion and cell biology in the future.

**Figure 4.**
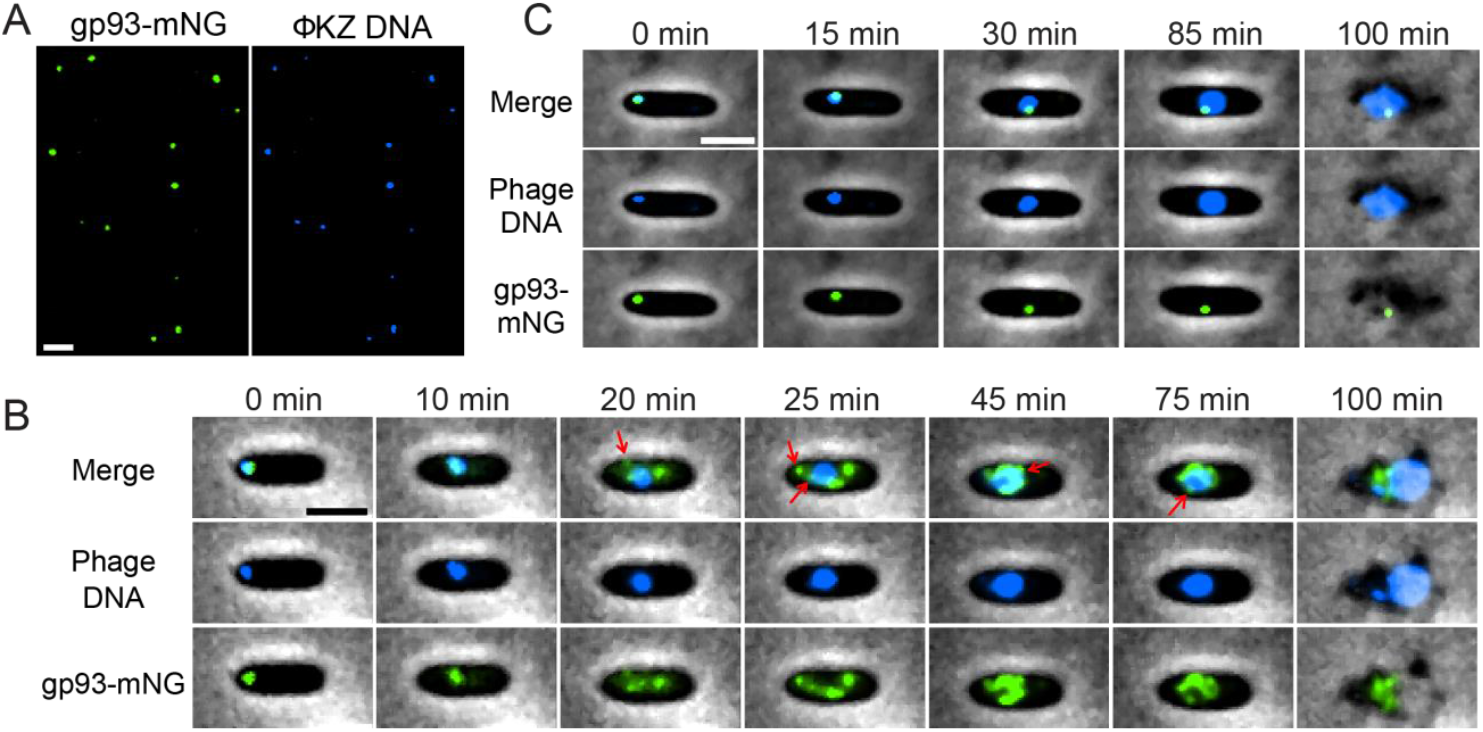
Chromosomal fluorescent labeling of an inner body protein of ΦKZ. (A) Visualization of individual phage particles under the fluorescence microscope. *orf39* encoding one of the major components of ΦKZ inner body is genetically fused to mNeonGreen. Each mutant phage particle is visible as a green focus (left), due to the packaging of gp93-mNeonGreen in the capsid. ΦKZ genomic DNA is labeled by DAPI (right). mNeonGreen and DAPI signals colocalize very well and individual virions are easily distinguishable. (B) Overlay images (phase-contrast and fluorescent channels) from a time-lapse movie depicting a representative PAO1 cell being infected by a gp93-mNeonGreen mutant phage. gp93-mNeonGreen and phage DNA are shown as green and blue signals, respectively. Red arrows point to newly assembled phage capsids, both originated on the cell inner membrane (20 min) and anchored on the phage nucleus surface for subsequent phage DNA packaging (25, 45, and 75 min). (C) Overlay images from a time-lapse movie showing a PAO1 cell being infected by a WT ΦKZ loaded with gp93-mNeonGreen fusions. Packaged gp93-mNeonGreen appears as a green focus and remains bound to phage DNA throughout the infection cycle. Scale bar denotes 2 μm.

To assess whether the deletions of *phuZ* or *orf93* impact growth in a strain-dependent manner, we challenged a panel of 21 *P. aeruginosa* clinical strains with the mutant phages. Plaque assays showed that host ranges of both mutants were quite similar to WT (Figure 3D, S3), suggesting that these knockouts, and Cas13a-mediated genetic engineering in general, has no impact on the ΦKZ host range.

The phage nucleus is primarily composed of gp54 [28]. We were unable to knockout or fluorescently label *orf54*, even when wild-type gp54 was provided by expressing from a plasmid *in trans*. The primers used to amplify the region of editing generated multiple bands for both deletion and tag-addition mutant variants (Figure S4A). N- or C-terminal fusion of gp54 with mCherry tags yielded similar results. Whole genome sequencing of an isolated *orf54 “pseudo* knock-out” strain revealed that part of the editing plasmid was integrated upstream of *orf54*, while the *orf54* gene was left intact (Figure S4B). A similar attempt to delete the structural gene (*orf146*) and a cluster of IB genes (*orf89-orf93*) also failed, while deletion of accessory genes *orf241* and *orf241-242* succeeded but yielded no obvious phenotype. These results highlighted that the CRISPR-Cas13a counter selection system is a strong and efficient phage genome engineering tool, but the modification of phage essential genes remains challenging.

### Precise genome engineering of clinical phage OMKO1

We next explored the versatility of our phage engineering platform by editing the genome of a clinical jumbo phage. We selected OMKO1, a *P. aeruginosa* phage with a ~280 kbp genome that has high sequence identity to ΦKZ (98.23%). OMKO1 can drive re-sensitization of surviving *P. aeruginosa* cells to small-molecule antibiotics. This phage has been used for phage therapy as emergency treatment for chronic infections caused by antibiotic resistant *P. aeruginosa* [34], and it is currently being tested in a phase I/II clinical trial (CYPHY, https://clinicaltrials.gov/ct2/show/NCT04684641). With this phage, we tested whether we could insert “DNA barcodes” without impacting host range, for downstream clinical applications. Insertion of a DNA tracking signature into clinical phages would enable differentiation from naturally occurring phages during the manufacturing process and following administration to patients.

Two engineered OMKO1 strains were generated, one with *acrVIA1* and a 120-nt barcode inserted downstream of the capsid gene, and another with *acrVIA1* integrated upstream of the shell gene (Figure 5A, Table 1). The presence of the desired inserts were confirmed with whole genome sequencing of both OMKO1 engineered strains, and no unintended genetic changes occurred. Moreover, both strains exhibited strong resistance to Cas13a targeting (Figure 5B), owing to the expression of *acrVIA1* from the phage genomes. The host range and virulence of the two engineered OMKO1 variants together with the parental phage was then assessed on 22 *P. aeruginosa* clinical strains (including PAO1). The experiment was performed in a microplate liquid assay, where phage variants were individually mixed with each host strain at a MOI~1 and MOI~0.01. All three phages displayed the same host range (Figure 5C, S5), capable of infecting and suppressing growth of 20/22 (91%) clinical strains tested. Infections at high MOI (MOI = 1) resulted in a broader host range and greater bacterial growth suppression, while low MOI (MOI = 0.01) infections were able to suppress cell growth of 12/22 (55%) hosts. All phages exhibited similar virulence across all hosts with small differences in 5/22 strains. In 2/22 hosts (FB-89 and FB-92), both engineered OMKO1 strains displayed significantly higher liquid assay scores than the parental phage (*p* < 0.05). While mutant OMKO1::Acr-BC had weaker virulence than the parental phage in 3/22 strains. Altogether, these results indicate that OMKO1’s host range was not affected, and virulence was impacted only modestly by inserting *acrVIA1* or *acrVIA1* and a barcode in the two selected genome locations under the tested conditions.

**Figure 5.**
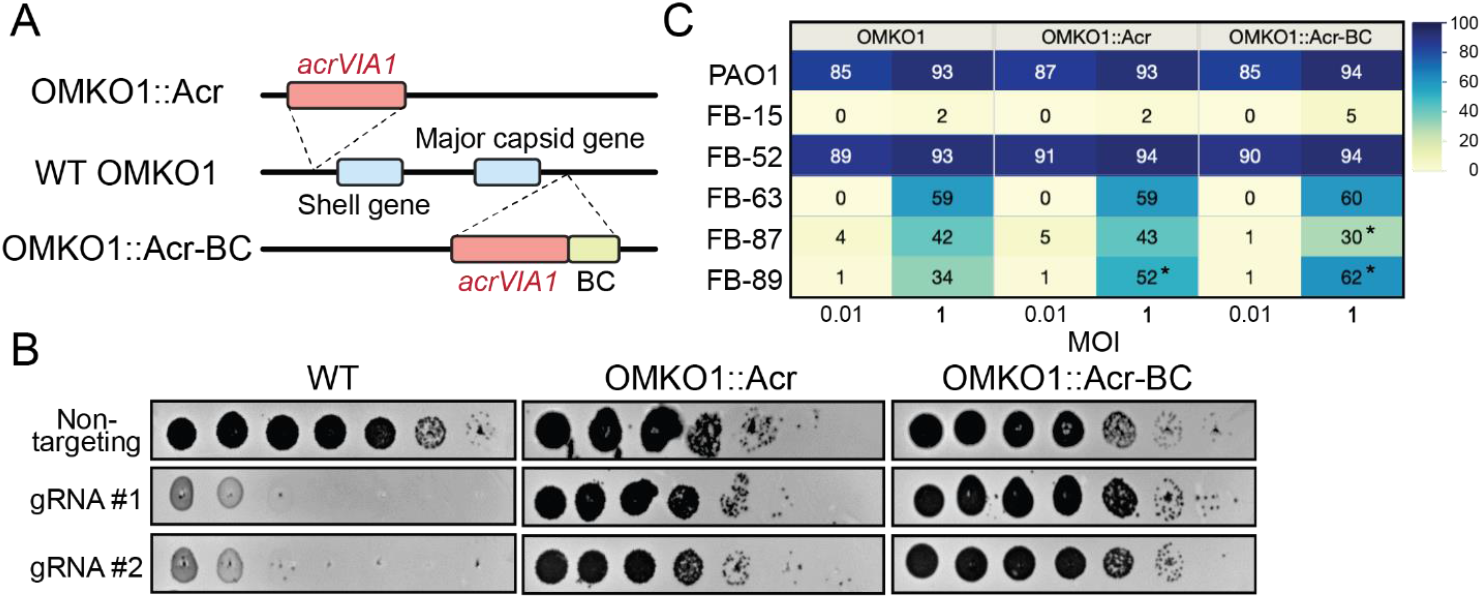
Genetic engineering of therapeutic jumbo phage OMKO1. (A) Schematic of genomes of two engineered OMKO1 variants where indicated gene fragments are integrated into the WT genome. OMKO1::Acr, *acrVIA1* is inserted upstream of the shell gene. OMKO1::Acr-BC, *acrVIA1* is inserted together with a barcode sequence (BC) downstream of the major capsid gene. (B) Plaque assays of WT OMKO1 and mutants. Both engineered variants exhibit robust resistance against distinct CRISPR-Cas13a crRNAs, suggesting that the incorporated AcrVIA1 is successfully functioning. (C) Determination of host range of WT OMKO1 and mutants on representative *P. aeruginosa* clinical strains by microplate liquid assay at MOI of 0.01 and 1. Data are presented as the mean liquid assay scores across three independent experiments. Asterisks (*) indicate significant difference between WT and mutants as determined by Students’ T-test (*p* < 0.05). The color intensity of each phage-host combination reflects the liquid assay score, with the darker color the stronger intensity displaying a greater score. (The full table of plaque assays is shown in Figure S5)

### Application of CRISPR-Cas13a phage engineering to a small lytic Podophage

To evaluate the applicability of the CRISPR-Cas13a-mediated genome editing approach to other virulent phages, we selected *P. aeruginosa* phage PaMx41. PaMx41 is a Podophage isolated from environmental and sewage water samples in Central Mexico [35]. Its genome is approximately 43.5 kbp long and harbors 55 open reading frames (ORFs), ~70% of which have unknown function [36]. Remarkably, we discovered that PaMx41 appears to be resistant to many DNA-targeting CRISPR-Cas systems (Type I-C, II-A, and V-A) and showed partial sensitivity (~10-fold reduction in efficiency of plating) to Type I-F to a degree that is not sufficient for counter selection (Figure 6A). In contrast, when the transcripts of the major capsid gene (*orf11*) were targeted by CRISPR-Cas13a, PaMx41 exhibited strong sensitivities to specific crRNAs (Figure 6B). Following the same approach as we developed to engineer ΦKZ, we successfully replaced one hypothetical gene (*orf24*) and its downstream non-coding region with *acrVIA1* and isolated a pure mutant strain using an efficient gRNA (#5) (Figure 6B, Table 1). The mutant showed expected anti-CRISPR activities against different *orf11*-targeting crRNAs, indicating that the incorporated *acrVIA1* was expressed and properly functioning (Figure 6C). No other obvious phenotype was observed for the PaMx41Δ*orf24* phage. Notably, initial PCR screening for recombinant plaques showed that 100% of surviving phages were desired recombinants, with no spontaneous escaper plaques. The data suggest that the RNA-targeting Cas13a system holds great promise for becoming a universal genetic editing tool to deal with previously intractable phages.

**Figure 6.**
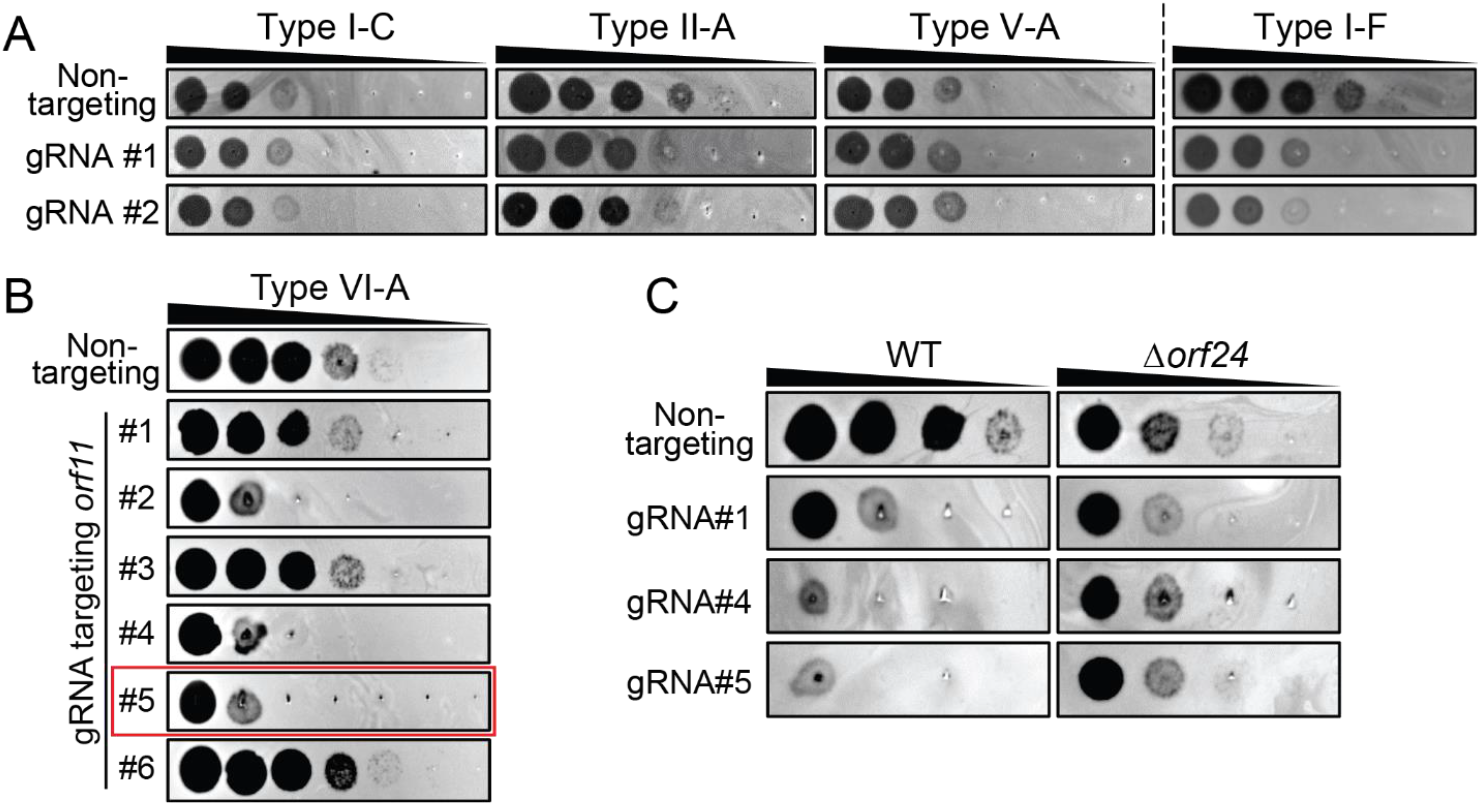
Genetic engineering of Podophage PaMx41. (A) Plaque assays showing PaMx41 resistance to a broad variety of DNA-targeting CRISPR-Cas systems. (B) PaMx41 exhibits significant sensitivity to CRISPR-Cas13a. gRNA#5 highlighted in the red frame has been used for PaMx41 genome engineering. (C) PaMx41 Δ*orf24* mutant strain is resistant to diverse crRNAs of CRISPR-Cas13a, due to the expression of AcrVIA1 from the phage genome.

## Discussion

We exploited the RNA-targeting CRISPR-Cas13a system in conjunction with homologous recombination to achieve genetic modification of jumbo phage ΦKZ, OMKO1, and phage PaMx41 that all resist DNA-targeting CRISPR-Cas systems. CRISPR-Cas13a-mediated counter-selection recovered rare (~10^−5^) phage recombinants from a large pool of wild-type phages. Many studies have uncovered that phages can hamper CRISPR-Cas activities, for example, by repressing transcription of endogenous CRISPR-Cas components [37, 38], possessing covalent DNA modifications [39–42], or encoding anti-CRISPR proteins (recently reviewed in [43]). Furthermore, the assembly of a proteinaceous nucleus-like structure that shields phage genomes from attack by distinct DNA-targeting nucleases [20, 21] represents the ultimate “anti-CRISPR/anti-RM” mechanism. Therefore, development of a phage genomic manipulation approaches that target a relatively constant and exposed molecule, mRNA, may provide a near-universal approach. Moreover, Cas13 is rarely encoded in bacteria and thus most phages are not expected to encode anti-Cas13a proteins.

Applying gene editing to this phage allowed us to query endogenous gene function and essentiality for the first time. We observed that Δ*phuZ* mutant phages mispositioned the phage nucleus during viral intracellular development. Previous studies revealed that newly assembled phage capsids trafficked along PhuZ filaments towards the phage nucleus for viral DNA packaging [29]. However, our work suggests that successful DNA loading into capsids is not dependent on PhuZ. Moreover, loss of PhuZ appeared not to affect burst size, in contrast with a previous report using over-expression of a catalytic mutant and microscopy to estimate burst size [30]. Overall, *phuZ* seems to be a *bona fide* nonessential gene for ΦKZ. The evolutionary advantage of encoding tubulin in this jumbo phage and many others requires further investigation. Furthermore, a chromosomal fluorescent label on gp93 demonstrated that it is packaged in the phage head, injected with the genome, and massively synthesized later in infection, with peri-nuclear localization. The labeling not only allows us to visualize individual virions under the microscope, but also to observe the injection of this inner body protein into the host cell, which had been previously suggested with little evidence [33, 44].

One major challenge of using Cas13a in our experience has been the wide variability of crRNA efficacy. Future studies focusing on the optimization of crRNA design for phage targeting or perhaps the implementation of other RNA-targeting enzymes, such as Cas13 orthologues or Cas7-11 [45, 46] will be important. However, we circumvent this problem by implementing an anti-CRISPR selectable marker [17] to ensure that the same strong guide can be used for all genetic manipulations. The downside is that this limits the user to a single perturbation, however, double and triple mutants are possible in principle if one uses crRNAs specific to the site of editing or removes the *acr* gene from the genome.

Altogether, the RNA-targeting CRISPR-Cas13a counter-selection tool should be applicable to a broad range of phages and enable downstream high throughput phage engineering. The ability to precisely and efficiently generate synthetic phages with desired features will not only benefit phage therapeutic applications but also advance our understanding of fundamental phage biology and phage-bacteria interactions.

## Supporting information

Genome Sequencing files

## Acknowledgements

J.B.-D. was supported by the National Institutes of Health (R01GM127489), the Vallee Foundation, the Searle Scholarship, the Innovative Genomics Institute, and the UCSF Program for Breakthrough Biomedical Research funded in part by the Sandler Foundation. This work was also supported by research funds from Felix Biotechnology.

We are grateful to Luciano Marraffini (The Rockefeller University) for providing the plasmid pAM383. We thank Paul Turner (Yale University) for providing phage OMKO1 and *P. aeruginosa* clinical isolates. We thank Gabriel Guarneros Peña at Centro de Investigación y de Estudios Avanzados for providing phage PaMx41. We thank Tomer Rotstein for his generous assistance with NGS data processing and interpretation. We thank members of the Bondy-Denomy laboratory for productive conversations and generous suggestions for our work.

## Materials and methods

### Strains, DNA oligonucleotides and plasmid constructions

All bacterial and phage strains, spacer sequences, and primers used in this study are listed in Tables S1, S2, and S3, respectively.

The crRNAs designed for CRISPR-Cas13 targeting were constructed in the pHERD30T backbone. The pHERD30T-crRNA Version 2 was constructed by thermal annealing of oligonucleotides oSDM465 and oSDM466 and phosphorylation by polynucleotide kinase (PNK). The annealed product was introduced by Gibson assembly into pHERD30T linearized by PCR using oligonucleotides oSDM457 and oSDM458. Proper construction of the expression vector was verified by Sanger sequencing. The pHERD30T-crRNA Version 3 was constructed just as for Version 2, but the crRNA-coding insert was instead composed of oligonucleotides oSDM455 and oSDM456. Both Version 2 and Version 3 of this plasmid were designed such that cleavage by BsaI would generate a linear plasmid that would accept annealed oligonucleotide spacers via ligation. Oligonucleotide pairs with repeat-specific overhangs encoding spacer sequences were annealed and phosphorylated using T4 polynucleotide kinase and then cloned into the BsaI-digested empty vectors. Cloning procedures were performed in commercial *E. coli* DH5α cells (New England Biolabs) according to the manufacturer’s protocols. The resulting crRNA plasmids were electroporated into *P. aeruginosa* PAO1 strain harboring the *tn7::cas13a^Lse^* (SDM084) on the chromosome as described previously [20]. Gene expression was induced by the addition of L-arabinose at a final concentration of 0.3% and isopropyl β-D-1-thiogalactopyranoside (IPTG) at a final concentration of 1 mM.

To construct template plasmids for homologous recombination, homology arms of >500 bp in length were amplified by PCR using the ΦKZ genomic DNA as the template. To prevent Cas13a cleavage, several synonymous mutations were introduced into the crRNA-targeting site of the left *orf120* homology arm by designing the reverse primer (JG064) to contain appropriate mismatches. The *acrVIA1* gene was amplified from plasmid pAM383 [24], a gift from Luciano Marraffini, The Rockefeller University. PCR products were purified and assembled as a recombineering substrate and then inserted into the NheI site of the pHERD30T backbone by Gibson Assembly (New England Biolabs) following the manufacturer’s protocols. The resulting plasmids were transformed into PAO1 by electroporation.

### Isolation of phage recombinants

Host strains bearing recombination plasmids were grown in LB supplemented with 10 mM MgSO_4_ and 50 μg/ml gentamicin, at 37°C with aeration at 250 rpm. When OD_600_ is around 2, Wild type ΦKZ was added into the culture at a MOI (multiplicity of infection) of 1 to allow infection to occur for ~18 hours. 2% volume of chloroform was added into the infection culture and left to shake gently on an orbital shaker at room temperature for 15 min, followed by centrifugation at 4,000 x g for 15 min to remove cell debris. The supernatant lysate was further treated with 2% of chloroform for 15 min and centrifuged again under the same conditions, followed by a 30-min treatment with DNase I (New England Biolabs) at 37°C. The resulting phage lysate containing both WT phages and recombinants are tittered on PAO1 strains bearing the CRISPR-Cas13a system with the most efficient crRNA (*orf120* guide#2) to screen for recombinants. Individual phage plaques were picked from top agar and purified for three rounds using the CRISPR counter-selection strain to ensure thorough removal of any remaining WT or escapers. Whether or not they are recombinant phages or Cas13a escaper phages were determined by PCR using appropriate pairs of primers amplifying the modified regions of the phage genome. Identified phages were further confirmed and analyzed by sequencing the PCR products or the whole genomes and then stored at 4°C.

### Phage plaque assay

Host strains were grown in LBM (LB supplemented with 10 mM MgSO_4_), 50 μg/ml gentamicin, 1 mM IPTG and 0.3% arabinose inducers for gene expression, at 37°C with aeration at 250 rpm for overnight. Phage spotting assays were performed using 1.5% LB agar plates and 0.42% LB top agar, both of which contained 10 mM MgSO_4_ and inducers. 100 μl of appropriate overnight culture was suspended in 3.5 ml of molten top agar and then poured onto an LB+10 mM MgSO_4_ agar plate, leading to the growth of a bacterial lawn. After 10-15 min at room temperature, 2 μl of ten-fold serial dilutions of phages was spotted onto the solidified top agar. Plates were incubated overnight at 37°C. Plate images were obtained using Gel Doc EZ Gel Documentation System (BioRad) and Image Lab (BioRad) software.

### Microplate Liquid Assay

Fresh overnight cultures were diluted to a cell concentration of 1×10^8^ cfu/ml in TSB media supplemented with 10mM MgSO_4_. Phage lysates were added to reach a MOI of ~1 and ~0.01 in a Corning Costar 96-well clear flat-bottom microplate (Thermo Fisher Scientific) sealed with a Breathe-Easy® sealing membrane (Merck KGaA). After the infection cultures were incubated at room temperature for 20 min, plates were incubated at 37°C, 800 rpm for 8 hours in a BioTek LogPhase 600 plate reader (Agilent Technologies, Inc.). Cell growth was monitored by measuring OD_600_ every 20 min. Each phage-host combination was performed in three biological replicates.

### Data Analysis

Growth curves for each phage-host combination were obtained by plotting OD_600_ after blank correction (baseline adjustment) against time. Each growth curve was transformed into a single numerical value by calculating the area under the curve (AUC) using the Trapezoid method. Then, AUCs were normalized as a percentage of the AUC of their corresponding uninfected control following the equation,

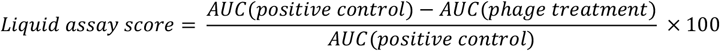

The resulting value, defined as the “liquid assay score”, represents how well the phage strain can repress the growth of a bacterial population over the course of the 8-hour experiment. No inhibition of bacterial growth would result in a liquid assay score of 0, and complete suppression would translate into a score of 100. Liquid assay scores were averaged using data from three biological replicates.

### Burst size measurement

Phage burst size was determined by one-step growth curve experiments. Briefly, the host PAO1 strain was grown in LB media to OD_600_ ~0.4. 1 ml of the cell culture was then centrifuged at 4,000 × g for 2 min and the cell pellet was resuspended in 50 μl of fresh LBM. Appropriate amount of phages was mixed with the cell culture to achieve an MOI of 0.01 to limit to single infections. The mixture was incubated on ice for 20 min for phage adsorption and transferred to a 37°C heat block for 10 min to trigger phage DNA injection. The infection mixture was centrifuged at 10,000 × g for 2 min. Transfer 10 μl of the supernatant into 990 μl of ice-cold SM buffer supplied with 2% chloroform. Titer to calculate the number of free phages. After discarding the supernatant to remove free phage particles, the pellet was resuspended in 1 ml of LBM, followed by incubation at 37°C with shaking at 250 rpm. Samples were collected at 10-min intervals until 90 min, and phage titer was determined immediately. Phage burst sizes were calculated by dividing the phage titers at ~50 min by the initial phage titers after subtracting free phages.

### Single-cell infection assay

1 ml of host cells was grown in LBM (LB supplemented with 10 mM MgSO_4_) and 50 μg/ml gentamicin (if necessary), at 37°C with aeration at 250 rpm for overnight. The overnight culture was diluted 1:100 into 5 ml of LBM and grown at 37°C with 250 rpm shaking until OD_600_ ~0.4. Next, 1 ml of cell culture was collected by centrifugation at 3,000 × g for 2 min at room temperature and concentrated by 25-fold in fresh LBM. 10 μl of cells were then mixed with 10 μl of appropriate phage strains to reach an appropriate MOI, followed by incubation at 30°C for 10 min to allow for phage infection. The infection mixture was further diluted by 10-fold into 50 μl of fresh LBM at room temperature. 1 μl of the diluted culture was gently placed onto a piece of agarose pad (~1 mm thick) with 1:5 diluted LBM, arabinose (0.8%), and DAPI (5 μg/ml; Invitrogen&#153, No. D1306). A coverslip (No.1.5, Fisher Scientific) was gently laid over the agarose pad and the sample was imaged under the fluorescence microscope at 30°C within a cage incubator to maintain temperature and humidity.

### Fluorescence microscopy and imaging

Microscopy was performed on an inverted epifluorescence (Ti2-E, Nikon, Tokyo, Japan) equipped with the Perfect Focus System (PFS) and a Photometrics Prime 95B 25-mm camera. Image acquisition and processing were performed using Nikon Elements AR software. During a time-lapse movie, the specimen was typically imaged at a time interval of 5 min at the focal plane for 2.5~3 h, through channels of phase contrast (200 ms exposure, for cell recognition), blue (DAPI, 200 ms exposure, for phage DNA), and green (GFP, 300 ms exposure, for Gp93-mNeonGreen).

### Next-generation sequencing (NGS)

To isolate phage genomic DNA, purified high titer lysates were treated with Benzonase Nuclease (Sigma) for 30 min at 37°C. Phage genomic DNA was extracted with a modified Wizard DNA Clean-Up kit (Promega) protocol. DNA samples were quantified with AccuGreen Broad Range dsDNA quantification kit (Biotium, USA) in a Qubit Fluorometer 2.0.

Purified phage genomic DNA was processed following Illumina DNA Preparation Protocol. Samples were sequenced on a MiSeq system (Illumina) with 300 cycles of paired-end sequencing, and loading concentration of 12 pM. Illumina short reads were downsampled to ~50-100× coverage and de novo assembled using SPAdes. The sequences of mutant phage strains were aligned to the reference genome in Geneious with the Mauve alignment algorithm to confirm the intended genomic edits.

The isolated *orf54 “pseudo* knock-out” phage strain (“*orf54*”) was sequenced using long-read sequencing. DNA samples were processed using SQK-LSK109 kit (Oxford Nanopore Technologies, UK). Libraries were sequenced using an R10.3 flow cell until the desired number of reads was achieved. Oxford Nanopore long reads were filtered for the longest high quality reads using Nanofilt and de novo assembled using Flye.

## Supplementary Information

**Figure S1.**
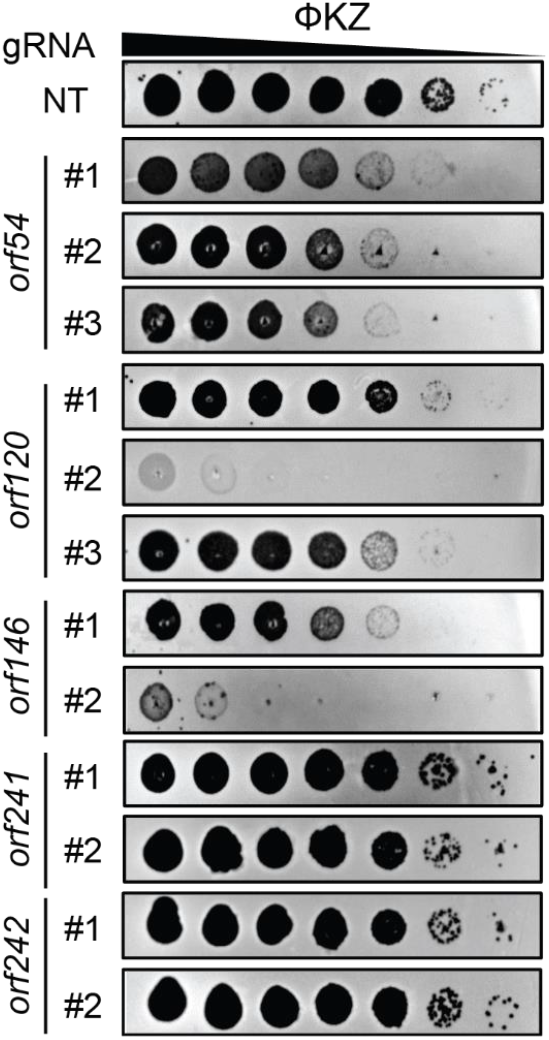
Plaque efficiency assays of distinct crRNAs of CRISPR-Cas13a targeting transcripts of diverse ΦKZ genes. Different crRNAs show varied degrees of Cas13a interference efficiency against ΦKZ. NT, non-targeting.

**Figure S2.**
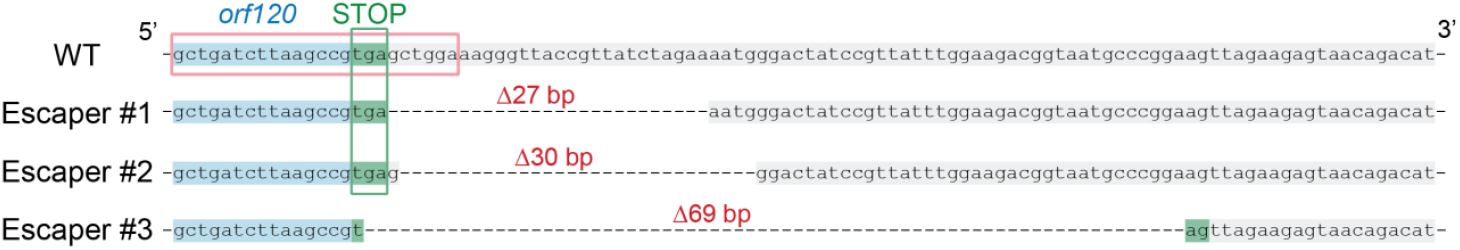
Sequence alignment of wild type ΦKZ and three escaper mutants at the engineered genomic site. Escaper mutants were isolated and verified by PCR and sequencing. The WT *orf120* sequence is highlighted in blue and the downstream region is highlighted in grey. The stop codon (TGA) of *orf120* is highlighted in green and Escaper #3 reconstitutes it to TAG. The sequence in the red frame matches the spacer sequence of the crRNA that was used to target and eliminate WT phages. Deletions were indicated by dashed lines and their corresponding numbers of absent base pairs.

**Figure S3.**
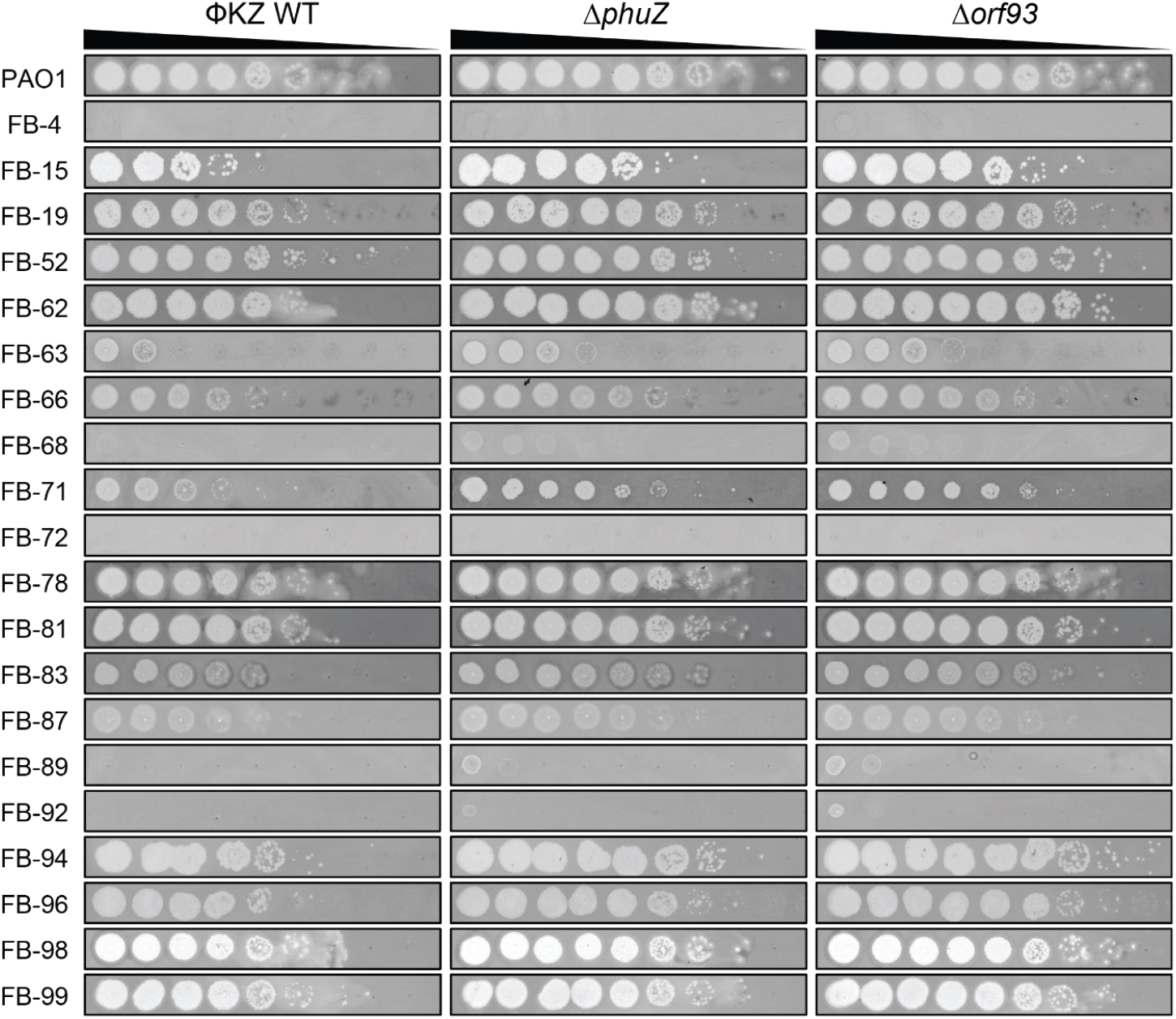
Determination of host range of ΦKZ Δ*phuZ* and Δ*orf93* mutants by plaque assay on *P. aeruginosa* clinical strains.

**Figure S4.**
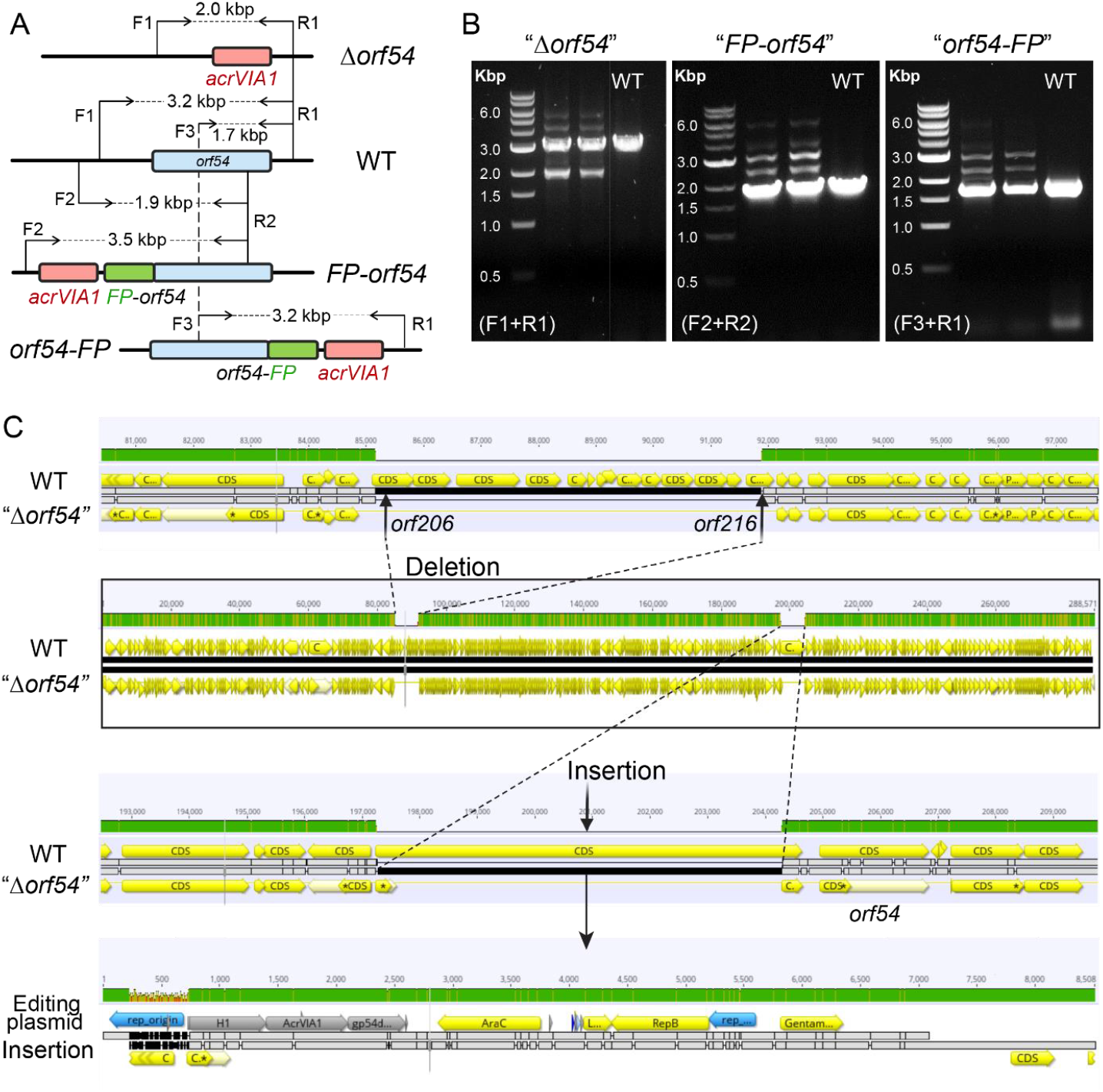
Failure of genetic editing the shell gene (*orf54*) in ΦKZ. (A) Schematic of genomes of WT ΦKZ and three mutated *orf54* variants, “Δ*orf54*”, “*FP-orf54*”, and *“orf54-FP”*, at the editing site. *orf54, acrVIA1*, and fluorescent protein (FP) are shown as blue, red, and green rectangles, respectively. F and R indicate forward and reverse primers, respectively, for PCR confirmation of *orf54* engineering. (B) PCR confirmation of the indicated *orf54* mutants using their corresponding pair of primers. All three mutants generated multiple bands, including a band in the same size as the single band produced by WT. (C) Genome alignment of WT phage with the isolated *orf54 “pseudo* knock-out” mutant (“Δ*orf54*”). A gene cluster of ~ 7 kbp (*orf206 - orf216*) was missing in the mutant, likely as a result of phage packaging capacity. The majority of the editing plasmid used to generate recombinants was at the editing site, leaving the *orf54* intact.

**Figure S5.**
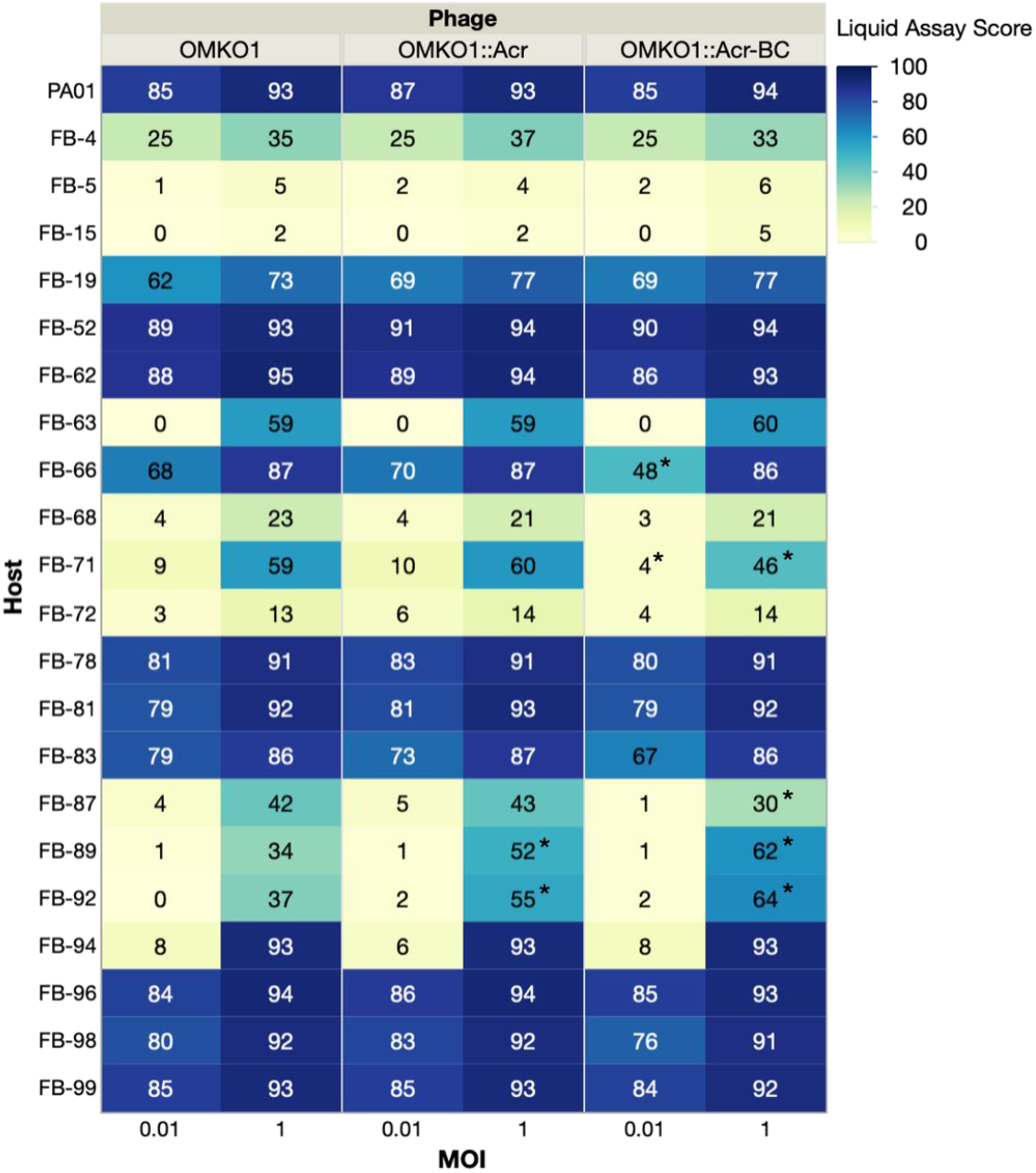
Host range assay of engineered OMKO1 variants. Host ranges were determined by microplate liquid assay at MOI of 0.01 and 1 on 22 *P. aeruginosa* clinical strains. The values are presented as the mean liquid assay scores across three independent experiments. Asterisks (*) indicate significant difference between WT and engineered strains as determined by Students’ T-test (*p* < 0.05). The color intensity of each phage-host combination reflects the liquid assay score, with the darker color the stronger intensity displaying a greater score.

**Movie S1. Time-lapse movie of a *P. aeruginosa* cell infected by a ΦKZ Δ*phuZ* mutant phage.**

Phage DNA is stained by DAPI and shown in blue.

**Movie S2. Time-lapse movie of a *P. aeruginosa* cell infected by a ΦKZ gp93-mNeonGreen mutant phage.**

Phage DNA is stained by DAPI and shown in blue. gp93-mNeonGreen is shown in green.

**Movie S3. Time-lapse movie of a *P. aeruginosa* cell infected by a WT ΦKZ packaged with gp93-mNeonGreen fusion proteins in the capsid.**

Phage DNA is stained by DAPI and shown in blue. gp93-mNeonGreen is shown in green.

**Table S1.** Bacterial strains and phages used in this work.

| Name | Description | Source/Reference |
| --- | --- | --- |
| **Bacterial strains** |  |  |
| PAO1 | *P. aeruginosa* | Lab collection |
| SDM084 | PAO1 *tn7::Lse cas13a* | [20] |
| BF# | *P. aeruginosa* CF clinical strains | Paul Turner Lab |
| **Phages** |  |  |
| ΦKZ |  | Alan Davidson Lab |
| JBD30 |  | Alan Davidson Lab |
| OMKO1 | Clinical *P. aeruginosa* phage | Paul Turner Lab |
| PaMx41 |  | Gabriel Guarneros Peña at Centro de Investigación y de Estudios Avanzados (KU884563) |

**Table S2.**
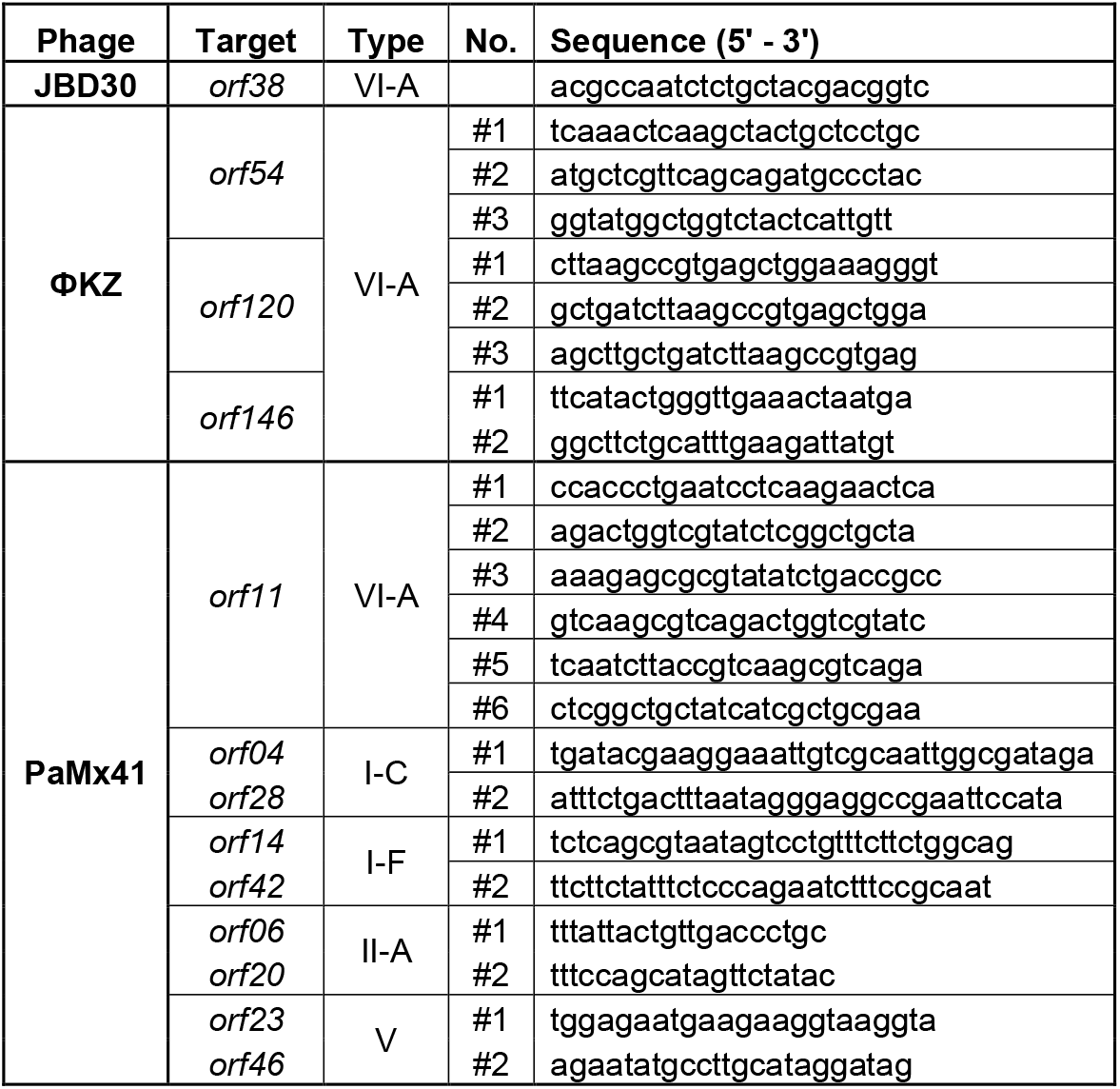
crRNA sequences.

**Table S3.**
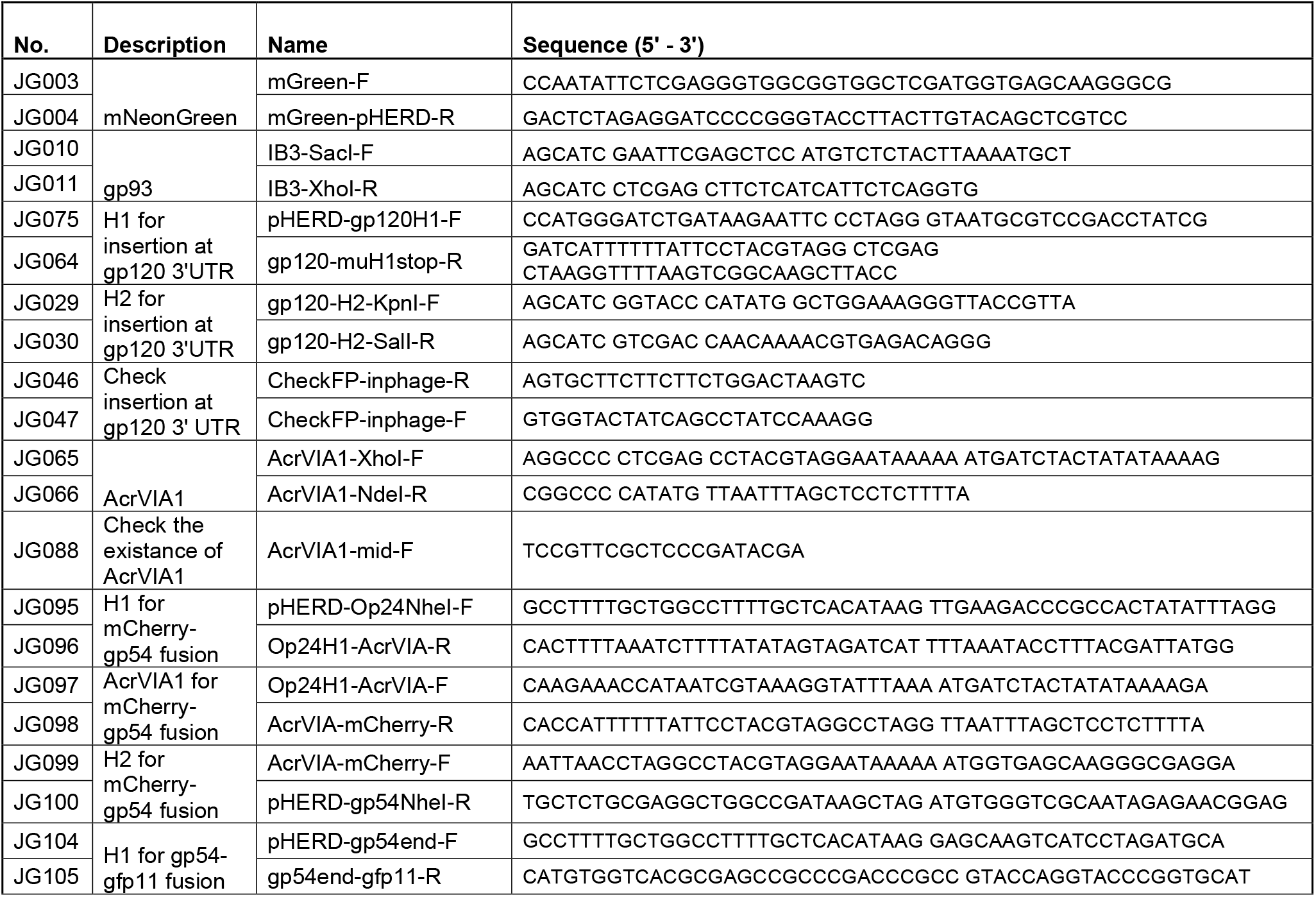
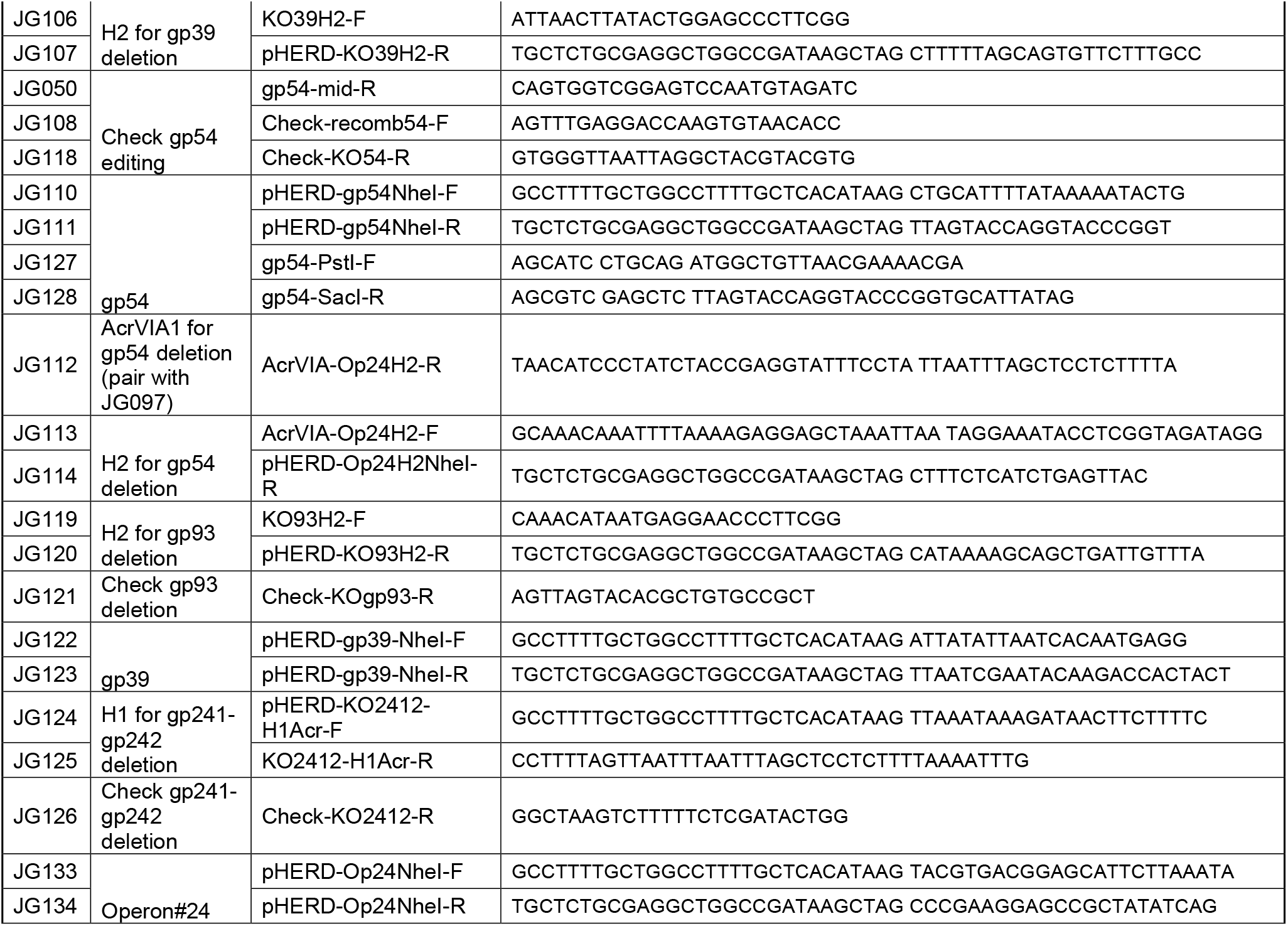
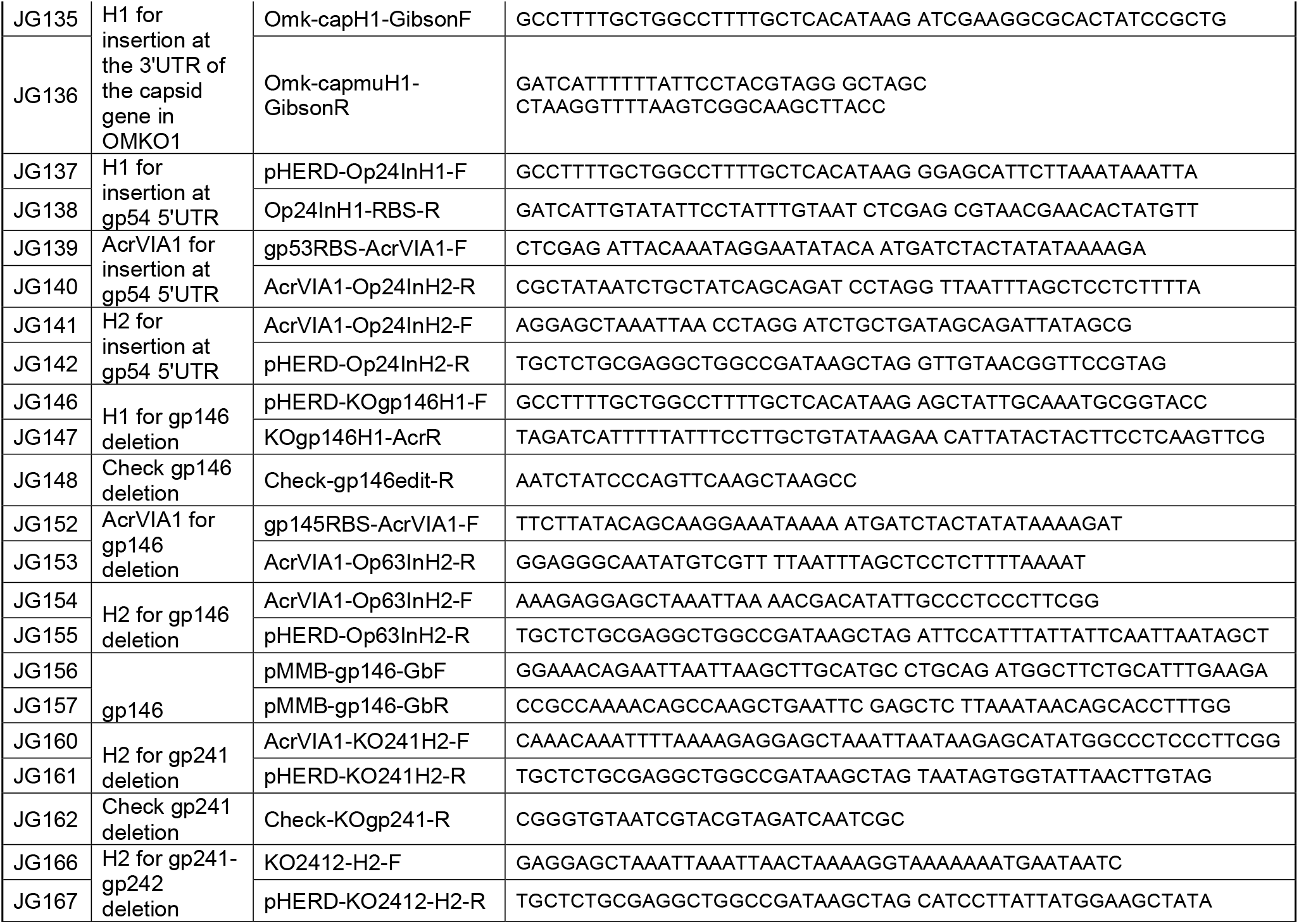
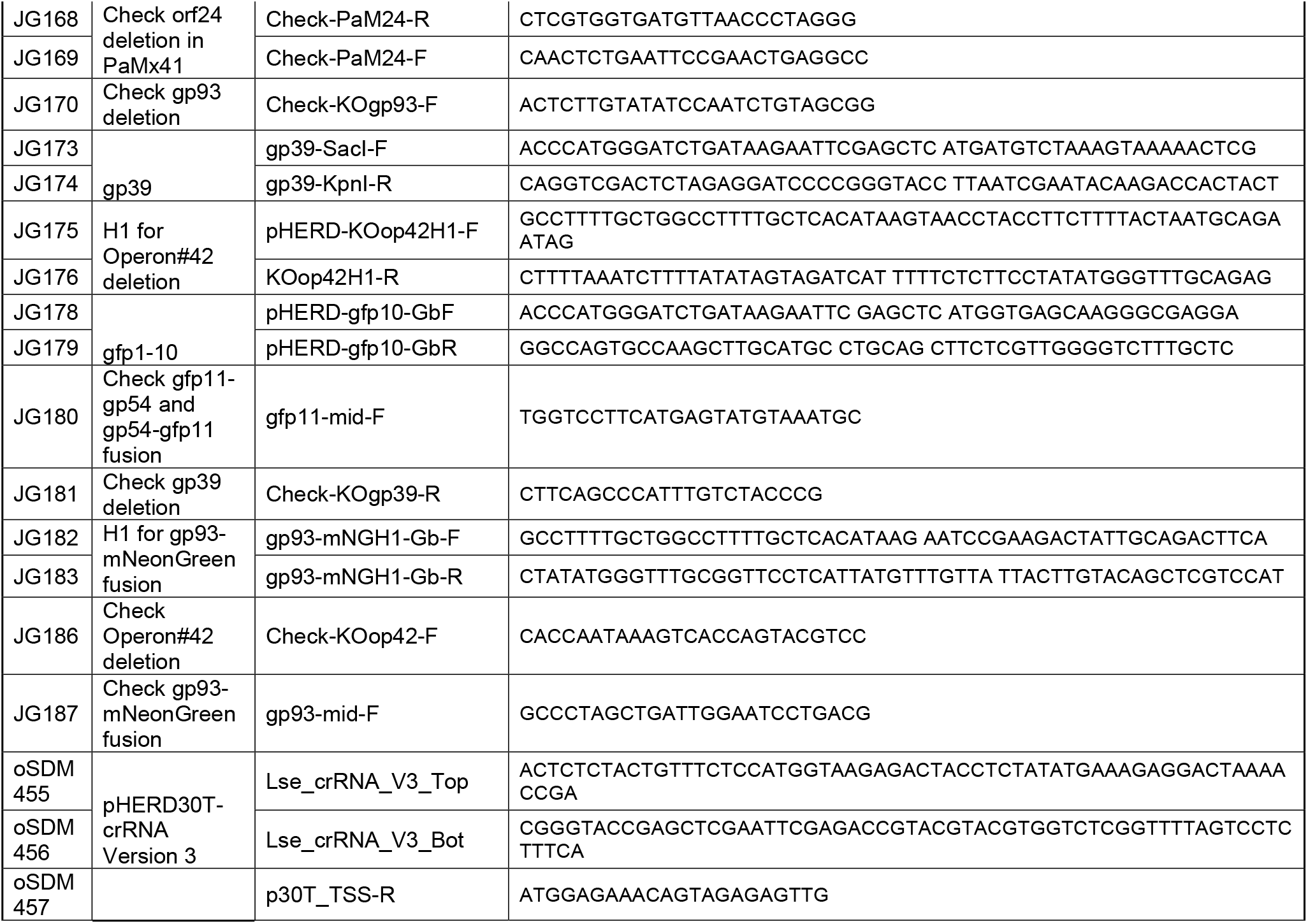
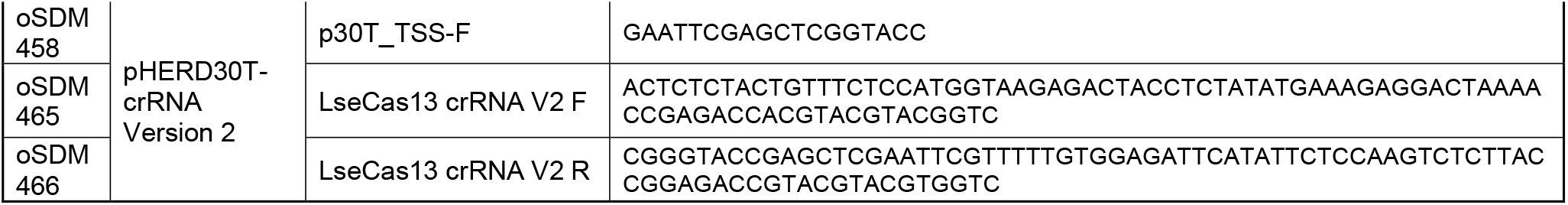
Primers.

